# LoVis4u: Locus Visualisation tool for comparative genomics

**DOI:** 10.1101/2024.09.11.612399

**Authors:** Artyom A. Egorov, Gemma C. Atkinson

**Affiliations:** Department of Experimental Medical Science, Lund University, Lund, Sweden

## Abstract

**Summary:** Comparative genomic analysis often involves visualisation of alignments of genomic loci. While several software tools are available for this task, ranging from Python and R libraries to standalone graphical user interfaces, there is lack of a tool that offers fast, automated usage and the production of publication-ready vector images.

Here we present LoVis4u, a command-line tool and Python API designed for highly customizable and fast visualisation of multiple genomic loci. LoVis4u generates vector images in PDF format based on annotation data from GenBank or GFF files. It is capable of visualising entire genomes of bacteriophages as well as plasmids and user-defined regions of longer prokaryotic genomes. Additionally, LoVis4u offers optional data processing steps to identify and highlight accessory and core genes in input sequences.

**Availability and Implementation:** LoVis4u is implemented in Python3 and runs on Linux and MacOS. The command-line interface covers most practical use cases, while the provided Python API allows usage within a Python program, integration into external tools, and additional customisation. Source code is available at the GitHub page: github.com/art-egorov/lovis4u. Detailed documentation that includes an example-driven guide is available from the software home page: art-egorov.github.io/lovis4u.

## Introduction

The exponential growth of microbial genome databases has unlocked numerous opportunities for comparative genomic analyses (1). Various tasks such as analysis of gene neighbourhood conservation (2,3), annotation of functional short ORFs (4,5), and investigation of genomic variability hotspots (6-8) often require visualisation of multiple genomic loci. Several software tools have been developed for this purpose. A subset of these have graphical user interfaces (GUIs), such as the Artemis Comparison Tool (9), Easyfig (10), GeneSpy (11), and Geneious Prime (geneious.com). Another category comprises web-based applications like Gene Graphics (12). Additionally, there are libraries such as the R packages genoPlotR (13) and gggenes (14), as well as the Python package GenomeDiagram (15). Some tools integrate multiple approaches, creating hybrid solutions. For example, GEnView is a Python pipeline combined with an interactive web application (16), and Clinker & clustermap.js (17) is a popular tool with a command-line interface and an interactive web application that can generate vector graphics. While many of these tools feature interactivity through GUIs or web applications, there is a lack of a user-friendly command-line tool suitable for handling multiple input genomes, flexible customisation options with batch mode possible, and with fast production of aesthetically pleasing publication-ready figures.

Here we present LoVis4u (**Lo**cus **Vis**ualisation), a scalable software tool designed for customisable and fast visualisation of multiple genomic loci. LoVis4u offers a command-line interface without requiring user-side scripting and provides a Python API for additional customization and integration within Python programs. In addition to visualisation features, our tool has optional data analysis steps that include protein clustering to find groups of protein homologues with subsequent identification of accessory and core genes that can be highlighted with visualisation. While LoVis4u was designed for annotating and visualising multiple bacteriophage genomes or their loci, it can be used for visualising defined regions of any prokaryotic genome.

## The LoVis4u workflow and features

The LoVis4u pipeline includes several default data processing steps, which are optional depending on user needs (Fig 1A). LoVis4u supports input data in either GenBank or extended GFF (concatenated with the corresponding nucleotide sequence in Fasta format) file format. GFF files in this format are produced by the widely used prokka (18) and pharokka (19) genome annotation tools.

**Figure 1:**
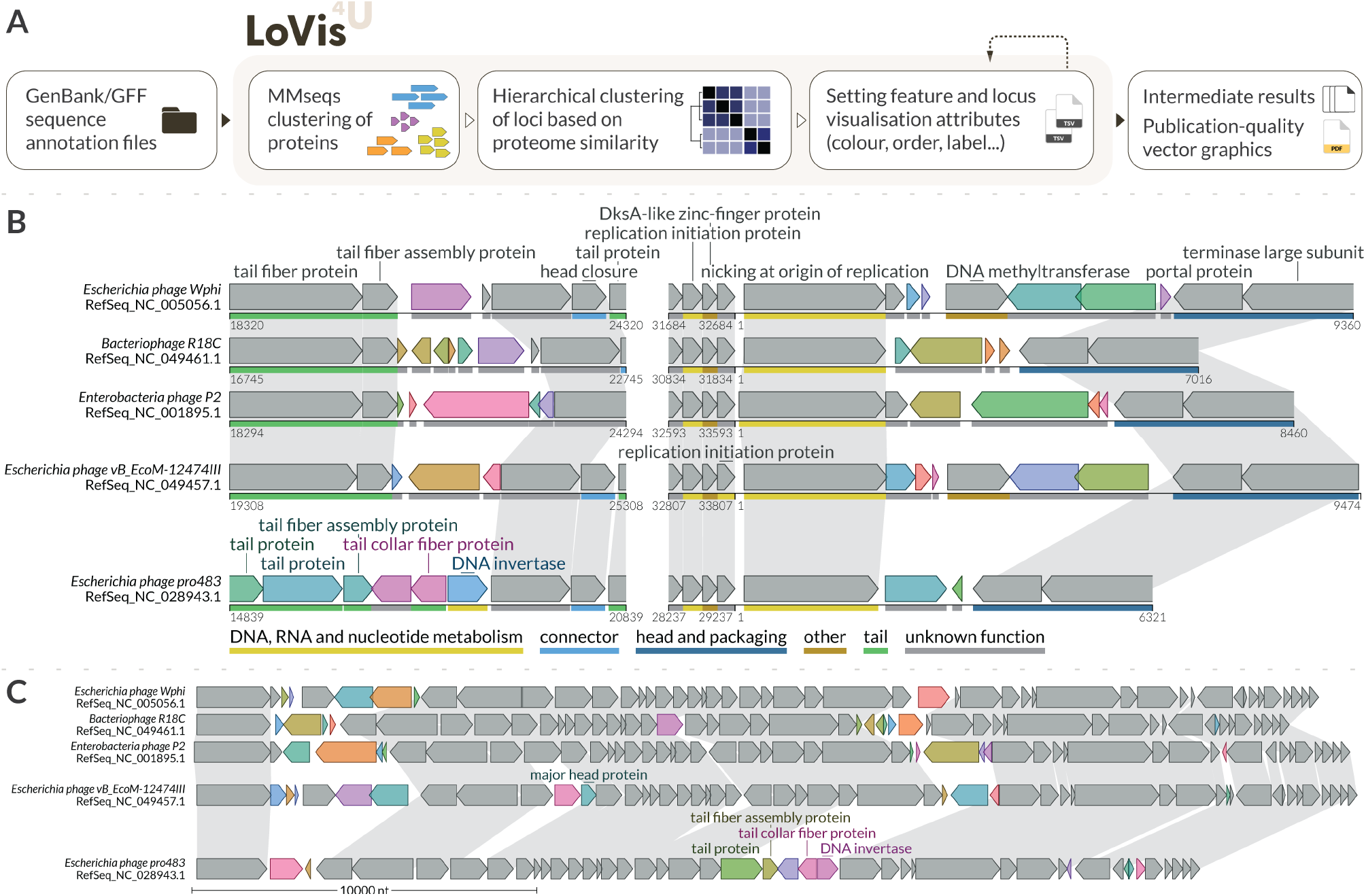
LoVis4u workflow and visualisation example. (A) Schematic description of the LoVis4u pipeline and its input and output (B) Visualisation of multiple regions for a set of P2-like phage genomes. Conserved genes are shown in grey while variable protein groups are highlighted with distinct colours. Homologous protein groups defined by MMSeqs2 clustering and located on different genomes are connected by grey homology lines. As can be set in the configuration file, protein labels are hidden for hypothetical proteins or proteins of unknown function. Labels for conserved proteins are shown only for the first occurrence since homology lines indicate additional occurrences. Label positions are arranged automatically by the algorithm to avoid overlaps of text. PHROG functional annotations (24) are indicated by coloured lines beneath each CDS, according to the colour code at the bottom of the panel. (C) Visualisation of a set of P2-like phage genomes that showcases a compact visualisation of full-length sequences, where functional annotation tracks and individual x-axes are hidden. Instead, a scale line at the bottom is displayed to indicate region size.

By default, LoVis4u applies the MMseqs2 (20) protein clustering algorithm to all encoded protein sequences to identify groups of homologous proteins. Alternatively, a table with predefined protein groups can be used as input. Based on the defined protein groups, LoVis4u constructs a matrix with pairwise proteome composition similarity scores that reflect the fraction of shared homologous proteins between sequences, and a corresponding proteome composition distance matrix as previously described (21). The clusters and order of input sequences for visualisation are determined using hierarchical clustering with average-linkage applied to the distance matrix. This approach allows us to consider a set of genes for each cluster analogously to pangenome analyses, where we define “*conserved”, “intermediate”* and “*variable”* protein group classes that are roughly equivalent to “*core”, “shell”*, and “*cloud”* terms of pangenomics (22). A similar clustering approach of sequences based on proteome equivalence was recently implemented in PhamClust (23). However, PhamClust does not include visualisation. A detailed workflow description is available on the LoVis4u home page (art-egorov.github.io/lovis4u).

LoVis4u can be run in quick-start mode, with few required options, but also has a range of advanced customisation options to give users full control over output. Sequence order, sequence clusters, and protein group classes can be manually specified through feature and locus annotation tables, which can be provided as optional arguments. Additionally, visualisation parameters such as figure width can be set with optional arguments or the configuration file. Users can also use these tables to customize colours and labels. A step-by-step guide on the homepage demonstrates how to experiment with and optimize visualisations using these features. Finally, LoVis4u uses the ReportLab library API to generate the output vector image, which is saved in PDF format. This format allows further editing of all objects in vector image-editing programs like Adobe Illustrator or Inkscape if needed.

Example LoVis4u output figures are shown in Fig 1(B-C), demonstrating a subset of the tool’s key features. The input data for these five P2-like phages includes genomes from the RefSeq database (25), along with optional specified visualisation parameters, including genomic coordinate limits. LoVis4u allows users to specify multiple regions for each sequence, which are displayed on a single line. Additionally, if a sequence is circular (as is the case with many phage and plasmid sequences), the visualisation will continue without a gap between sequence end and start coordinates. By default, LoVis4u highlights “variable” proteins by automatically assigning different colours to each homologous protein group. Alternatively, conserved proteins can be highlighted. LoVis4u also automatically interprets PHROG (24) functional group annotations, as provided in GFF and GenBank files produced by pharokka, and can display these annotations with a functional category line beneath the ORFs. LoVis4u is also suitable for full-length visualisation with a more minimalistic and compact design as shown in panel Fig. 1C. A step-by-step guide for creating these figures can be found in the user guide on the tool’s homepage.

To demonstrate full-length sequence visualisation, we have used LoVis4u to create a visualisation of the entire Basel collection of 78 phages (26) in a single figure, highlighting the variable genes (Supplementary File 1). The complete pipeline, which includes clustering of 13,630 protein sequences, ordering of the sequences by hierarchical clustering, and generating the graphic output, took only 50 seconds to run on an M1 MacBook Pro laptop.

## Implementation

LoVis4u is implemented in Python3 and uses multiple python3 libraries: biopython (27), bcbio-gff, scipy (28), configs, argparse, pandas (29), distinctipy (30), matplotlib (31), seaborn (32), and reportlab. LoVis4u also uses MMseqs2 (20) as a non-python dependency, which is embedded in the library.

The python LoVis4u package is available in PyPI (*python3 -m pip install lovis4u*), and the source code is provided on the GitHub page (github.com/art-egorov/lovis4u). Detailed documentation with an installation guide, and an example-driven manual are available on the LoVis4u home page (art-egorov.github.io/lovis4u).

To facilitate the use of LoVis4u in standard scenarios, we provide a command-line interface that that does not requires user-side scripting. LoVis4u is highly customisable with the main options accessible directly through the command-line interface. Additional parameters, such as colour palette, font typeface, and other preferences, can be defined in the configuration files. Additionally, LoVis4u offers a Python API, allowing further customization and seamless integration into Python programs.

## Conclusion

Comparative genomic analysis in microbiological studies often requires specialised visualisation tools for various tasks. Here, we present LoVis4u, a software tool designed to produce publication-quality figures of genomic loci with several automated analysis steps integrated into the pipeline. LoVis4u offers a much needed compromise between tools with advanced R/Python APIs and those with user-friendly graphical interfaces and interactivity. It can be used for fast and automated generation of multiple figures in bioinformatics pipelines or other libraries, or can be used as a stand-alone tool for data exploration. By sensitively finding and annotating conserved and variable regions, LoVis4u can facilitate comparative evolutionary analyses of genomes or genomic regions, and the discovery of new biology.

## Conflict of interest

The authors declare that there is no conflict of interest.

## Funding

The work was supported by grants to GCA from the Knut and Alice Wallenberg Foundation (number 2020-0037), Vetenskapsrådet (the Swedish research council; numbers 2023-02353 and 2022-01205) and eSSENCE@LU (number 10:2). The tool was tested using resources provided by LUNARC, The Centre for Scientific and Technical Computing at Lund University.

## Notes

### Competing Interest Statement

The authors have declared no competing interest.

## References

1. Koonin, E.V., Makarova, K.S. and Wolf, Y.I. (2021) Evolution of Microbial Genomics: Conceptual Shifts over a Quarter Century. Trends Microbiol, 29, 582–592.

2. Saha, C.K., Sanches Pires, R., Brolin, H., Delannoy, M. and Atkinson, G.C. (2021) FlaGs and webFlaGs: discovering novel biology through the analysis of gene neighbourhood conservation. Bioinformatics, 37, 1312–1314.

3. Pereira, J. (2021) GCsnap: Interactive Snapshots for the Comparison of Protein-Coding Genomic Contexts. J Mol Biol, 433, 166943.

4. Andrews, S.J. and Rothnagel, J.A. (2014) Emerging evidence for functional peptides encoded by short open reading frames. Nat Rev Genet, 15, 193–204.

5. Egorov, A.A. and Atkinson, G.C. (2023) uORF4u: a tool for annotation of conserved upstream open reading frames. Bioinformatics, 39.

6. Yutin, N., Tolstoy, I., Mutz, P., Wolf, Y.I., Krupovic, M. and Koonin, E.V. (2024) Jumping DNA polymerases in bacteriophages. bioRxiv.

7. Rousset, F., Depardieu, F., Miele, S., Dowding, J., Laval, A.L., Lieberman, E., Garry, D., Rocha, E.P.C., Bernheim, A. and Bikard, D. (2022) Phages and their satellites encode hotspots of antiviral systems. Cell Host Microbe, 30, 740–753 e745.

8. Hochhauser, D., Millman, A. and Sorek, R. (2023) The defense island repertoire of the Escherichia coli pan-genome. PLoS Genet, 19, e1010694.

9. Carver, T.J., Rutherford, K.M., Berriman, M., Rajandream, M.A., Barrell, B.G. and Parkhill, J. (2005) ACT: the Artemis Comparison Tool. Bioinformatics, 21, 3422–3423.

10. Sullivan, M.J., Petty, N.K. and Beatson, S.A. (2011) Easyfig: a genome comparison visualizer. Bioinformatics, 27, 1009–1010.

11. Garcia, P.S., Jauffrit, F., Grangeasse, C. and Brochier-Armanet, C. (2019) GeneSpy, a user-friendly and flexible genomic context visualizer. Bioinformatics, 35, 329–331.

12. Harrison, K.J., Crecy-Lagard, V. and Zallot, R. (2018) Gene Graphics: a genomic neighborhood data visualization web application. Bioinformatics, 34, 1406–1408.

13. Guy, L., Kultima, J.R. and Andersson, S.G. (2010) genoPlotR: comparative gene and genome visualization in R. Bioinformatics, 26, 2334–2335.

14. Wilkins, D. (2023).

15. Pritchard, L., White, J.A., Birch, P.R. and Toth, I.K. (2006) GenomeDiagram: a python package for the visualization of large-scale genomic data. Bioinformatics, 22, 616–617.

16. Ebmeyer, S., Coertze, R.D., Berglund, F., Kristiansson, E. and Larsson, D.G.J. (2022) GEnView: a gene-centric, phylogeny-based comparative genomics pipeline for bacterial genomes and plasmids. Bioinformatics, 38, 1727–1728.

17. Gilchrist, C.L.M. and Chooi, Y.H. (2021) clinker & clustermap.js: automatic generation of gene cluster comparison figures. Bioinformatics, 37, 2473–2475.

18. Seemann, T. (2014) Prokka: rapid prokaryotic genome annotation. Bioinformatics, 30, 2068–2069.

19. Bouras, G., Nepal, R., Houtak, G., Psaltis, A.J., Wormald, P.J. and Vreugde, S. (2023) Pharokka: a fast scalable bacteriophage annotation tool. Bioinformatics, 39.

20. Steinegger, M. and Soding, J. (2017) MMseqs2 enables sensitive protein sequence searching for the analysis of massive data sets. Nat Biotechnol, 35, 1026–1028.

21. Gapinska, M., Zajko, W., Skowronek, K., Figiel, M., Krawczyk, P.S., Egorov, A.A., Dziembowski, A., Johansson, M.J.O. and Nowotny, M. (2024) Structure-functional characterization of Lactococcus AbiA phage defense system. Nucleic Acids Res, 52, 4723–4738.

22. Brockhurst, M.A., Harrison, E., Hall, J.P.J., Richards, T., McNally, A. and MacLean, C. (2019) The Ecology and Evolution of Pangenomes. Curr Biol, 29, R1094–R1103.

23. Gauthier, C.H. and Hatfull, G.F. (2023) PhamClust: a phage genome clustering tool using proteomic equivalence. mSystems, 8, e0044323.

24. Terzian, P., Olo Ndela, E., Galiez, C., Lossouarn, J., Perez Bucio, R.E., Mom, R., Toussaint, A., Petit, M.A. and Enault, F. (2021) PHROG: families of prokaryotic virus proteins clustered using remote homology. NAR Genom Bioinform, 3, lqab067.

25. O’Leary, N.A., Wright, M.W., Brister, J.R., Ciufo, S., Haddad, D., McVeigh, R., Rajput, B., Robbertse, B., Smith-White, B., Ako-Adjei, D. et al. (2016) Reference sequence (RefSeq) database at NCBI: current status, taxonomic expansion, and functional annotation. Nucleic Acids Res, 44, D733–745.

26. Maffei, E., Shaidullina, A., Burkolter, M., Heyer, Y., Estermann, F., Druelle, V., Sauer, P., Willi, L., Michaelis, S., Hilbi, H. et al. (2021) Systematic exploration of Escherichia coli phage-host interactions with the BASEL phage collection. PLoS Biol, 19, e3001424.

27. Cock, P.J., Antao, T., Chang, J.T., Chapman, B.A., Cox, C.J., Dalke, A., Friedberg, I., Hamelryck, T., Kauff, F., Wilczynski, B. et al. (2009) Biopython: freely available Python tools for computational molecular biology and bioinformatics. Bioinformatics, 25, 1422–1423.

28. Virtanen, P., Gommers, R., Oliphant, T.E., Haberland, M., Reddy, T., Cournapeau, D., Burovski, E., Peterson, P., Weckesser, W., Bright, J. et al. (2020) SciPy 1.0: fundamental algorithms for scientific computing in Python. Nat Methods, 17, 261–272.

29. team, T.p.d. (2024). Zenodo.

30. Jack Roberts, J.C., Kian-Meng Ang, & Yannick Brandt. (2024). Zenodo.

31. Hunter, J.D. (2007) Matplotlib: A 2D Graphics Environment. Computing in Science & Engineering, 9, 90–95.

32. Waskom, M. (2021) seaborn: statistical data visualization. Journal of Open Source Software, 6.

