## Supplementary figures and images for "LoVis4u: Locus Visualisation tool for comparative genomics"

### Supplementary File 1

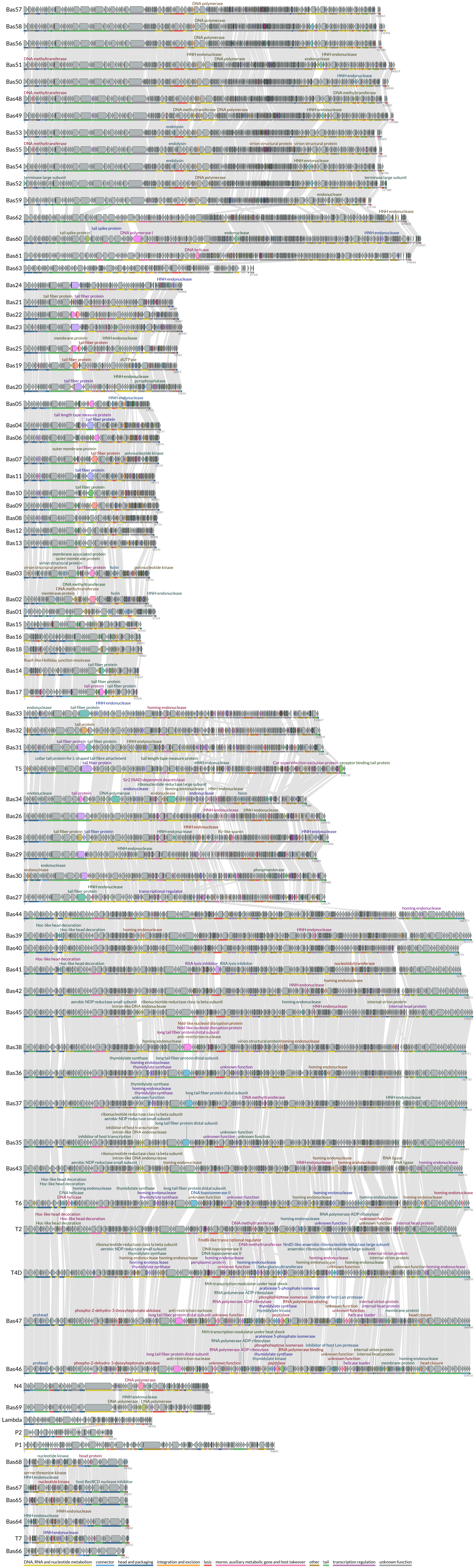
